## Supplementary Figures for "A prefrontal motor circuit initiates persistent movement"

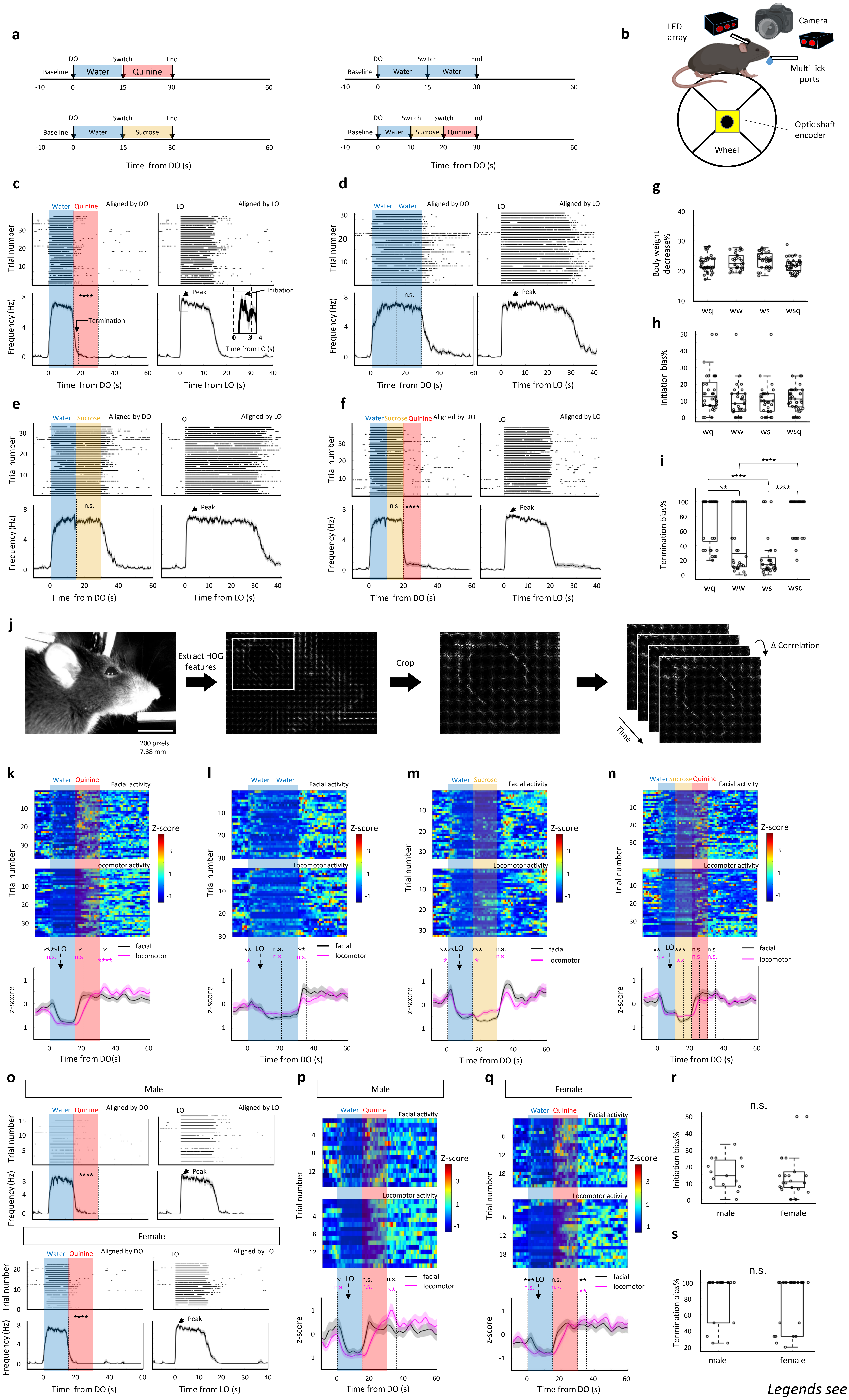


*Legends see next page*

**Supplementary Fig 1. Behavioral performance in persistent licking tasks**

**a.** Schematic of timeline for licking tasks per trial. **b.** Schematic of behavioral setup and equipment. **c-f.** Lick behavior relative to delivery onset (DO, left) and to $1^{st}$ lick onset (LO, right) after DO in four sessions. Top, raster plots showing lick behavior in the session water (w, 15s)-quinine (q, 15s) (c), water (w, 15s)-water (w, 15s) (d), water (w, 15s)-sucrose (s, 15s) (e), and water (w, 10s)-sucrose (s, 10s)-quinine (q, 10s). Bottom, lick frequency plot. The 'peak' indicates the maximum lick frequency. Wilcoxon signed-rank test, in comparison with baseline: ****p<0.0001, n.s. not significant. Values are mean ± s.e.m. **g.** Comparison of the percentage of body weight decrease among four sessions using one-way anova. Wq: n=37; ww: n=30; ws: n=33; wsq: n=39. **h & i.** Percentage of bias that started (h) or stopped (i) persistent lick (Methods). The trial numbers are the same as (g). One-way anova: p=0.1147 for initiation bias; p<0.0001 for termination bias. Bonferroni multi-compare: **p<0.01 ****p<0.0001. **j.** Illustration of calculating facial activity as 1-Δcorrelation (see Methods). **k-n.** Color-coded plot and traces showing facial and locomotor activity relative to water DO. We compared the epochs from DO to LO, from switch onsets to switch onsets+5s, from end onsets to end onsets+5s with the baseline. Wilcoxon signed-rank test: *p<0.05, **p<0.01, ***p<0.001, ****p<0.0001, n.s. not significant. Values are mean ± s.e.m. **o-s.** Comparison of licking performance (o), facial and locomotor activity (p and q), initiation bias (r), and termination bias (s) between male and female mice in w-q session. n_male=15; n_female=22. For all boxplot, the minima, maxima, and center bounds of box denote 25 percentile, median, and 75 percentile of data, respectively. The upper bound of whisker denote the highest data point, which lower than the sum of maxima bound and 1.5 times of box length. The lower bound of whisker denote the lowest data point, which higher than the subtraction of minima bound and 1.5 times of box length. *p<0.05, **p<0.01, ***p<0.001, ****p<0.0001, n.s. not significant. Values are mean ± s.e.m.

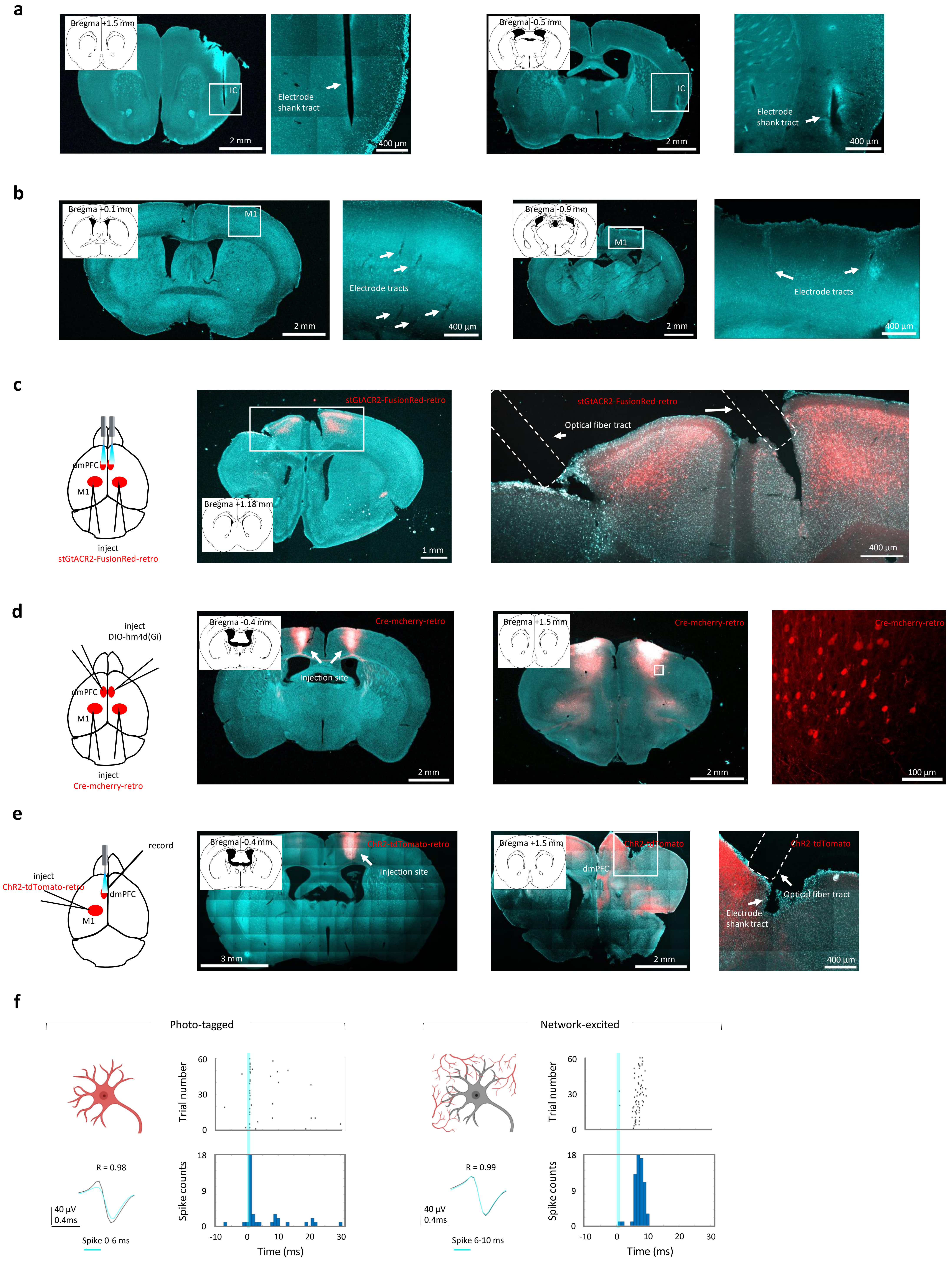


**Supplementary Fig 2. Histological and electrophysiological verification**

**a.** Histological verification of IC implant sites. From left to right: representative images of anterior IC implant is shown in 1st to 2nd columns and posterior IC implant is shown in 3rd to 4th columns. The magnified images are shown2nd and 4th columns. **b.** Histological verification of M1 implant sites. From left to right: representative anterior M1implant is shown in 1st to 2nd columns and posterior M1 implant is shown in 3rd to 4th columns. The magnified images are shown in 2nd and 4th columns. **c.** Histological verification of optic fiber implans for optogenetic silencing in bilateral dmPFC. **d.** Histological verification of viral injections for chemogenetic inhibition of hM4D labeled neuron in bilateral dmPFC. **e.** Histological verification of viral injections and electrodes and optic fiber implants for optogenetic identification of ChR2 labeled neuron in unilateral dmPFC. **f.** Electrophysiological verification of photo-tagged and network-excited neurons (see Methods).

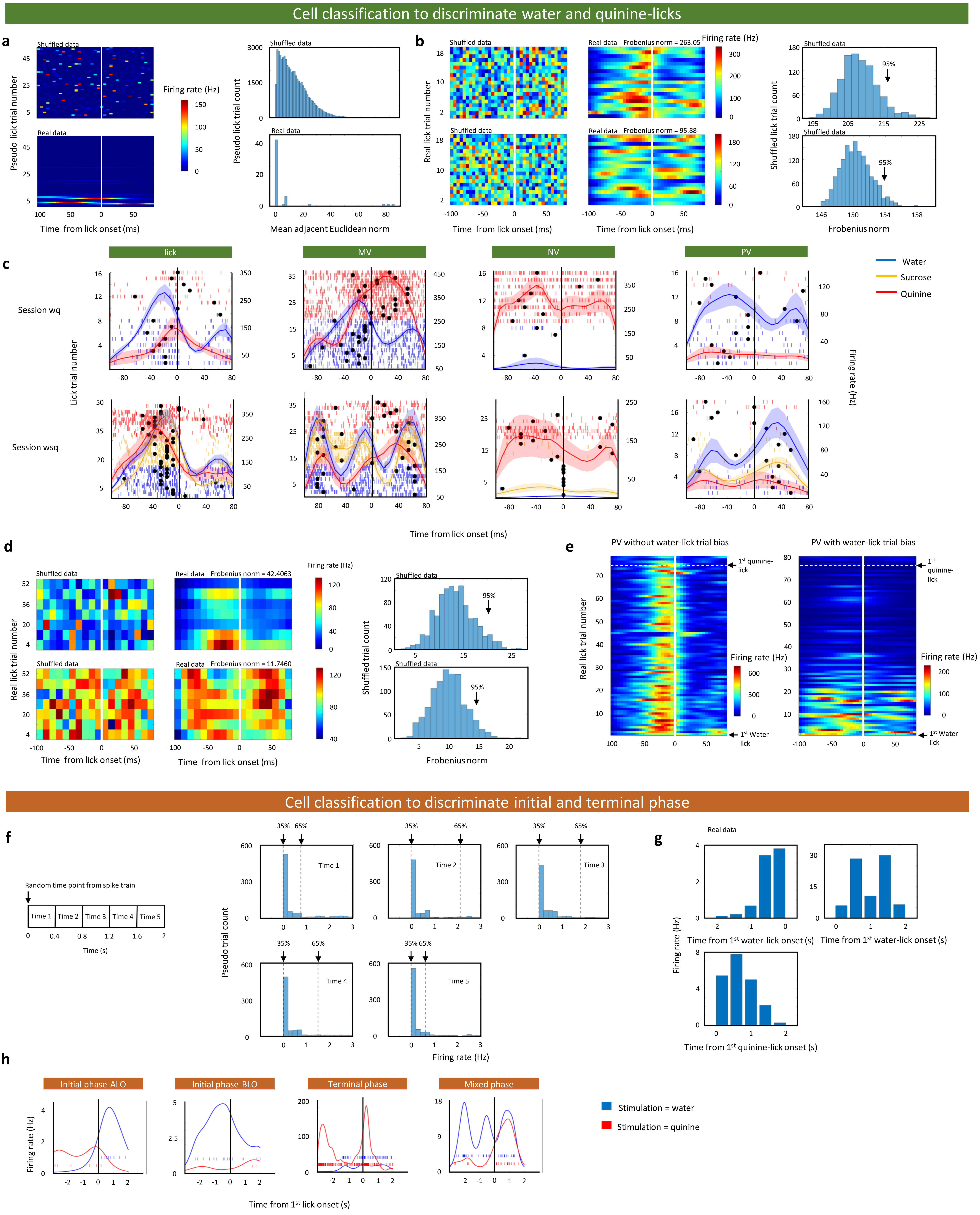


*Legends see next page*

**Supplementary Fig 3. Analysis of single-unit classification**

**a.** Procedure for shuffling baseline data. Left: color coded firing rates in the real and pseudo lick trials (the pseudo lick number equals to the real lick number after the baseline). Right: distributions of mean adjacent Euclidean norm obtained from the 1000 pseudo datasets and 1 real dataset. **b.** Procedure for categorizing the single-units with or without time bias. Top: example of a single-unit with time bias. Bottom: example of a single-unit without time bias. Left: color-coded firing rates showing shuffled data during lick trials. Middle: real data firing rates. Right: distributions of Frobenius norms of shuffled data. The single-unit was categorized as time bias when its Frobenius norm higher than 95 percentile of shuffled data. **c.** Spike raster and firing rates plots showing representative classified single-units that encode lick, mix valence (MV), negative valence (NV), and positive valence (PV) in the session water-quinine (wq, top) and the session water-sucrose-quinine (wsq, bottom). Black dots denote the peak firing rate at the each lick trial. **d.** Procedure for categorizing the neural representations with or without water-lick trial bias. Example of a single-unit with (top) or without (bottom) lick trial bias (matrix was binned with each 8 trials). Left: color-coded firing rates showing shuffled data during water-lick trials. Middle: real data firing rates. Right: distributions of Frobenius norms of shuffled data. The single-unit was categorized as water-lick bias when its Frobenius norm higher than 95 percentile of shuffled data. **e.** Color-coded firing rates showing the two examples of PV neural representations with (right) or without (left) water-lick trial bias. **f.** Procedure for generating pseudo data. Left: illustration of pseudo data selecting. Right: distributions of firing rates in 5 pseudo time points. **g.** Example of a single-unit firing rate in three recording epochs. **h.** Spike raster and firing rates plots showing representative classified single-units that represented initial phase-after lick onset (ALO), initial phase-before lick onset (BLO), terminal phase, and mixed phase.

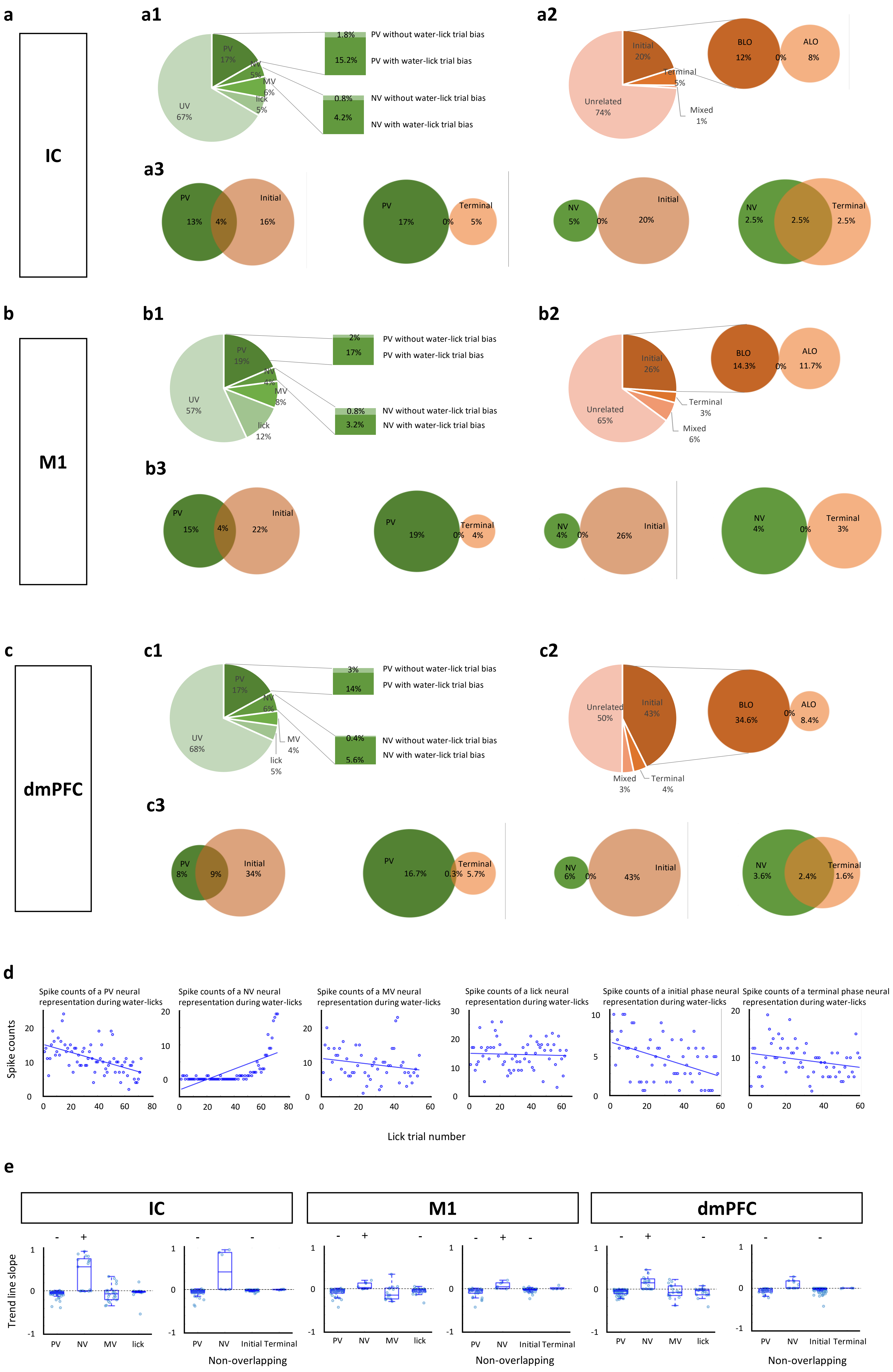


**Supplementary Fig 4. The percentage and firing trend of specific neural representations in the brain regions IC, M1, and dmPFC**

**a-c.** Percentage of the classified neural representations in IC (a), M1 (b), and dmPFC (c). **a1** & **b1** & **c1**: percentage of PV, NV, MV, lick, and UV (unrelated valence) neural representations. **a2** & **b2** & **c2**: percentage of initial phase (including ALO and BLO), terminal phase, mixed phase, and unrelated movement phase neural representations. **a3** & **b3** & **c3**: venn diagram showing the overlap and non-overlap percentage of indicated neural representations. **d.** The spike counts of indicated neural representations during water-licks. A blue dot denotes the spike times in the small scale window (LO-100ms to LO+80ms) of a water-lick trial. Blue lines represent the trend line of blue dots. **e.** Slopes of trend line of overall non-overlapping neural representation groups in IC, M1, and dmPFC as indicated. The symbol + and – represent the significant positive and negative value of its labeled trend line slope, respectively. + or - P<0.05, one sample t-test.

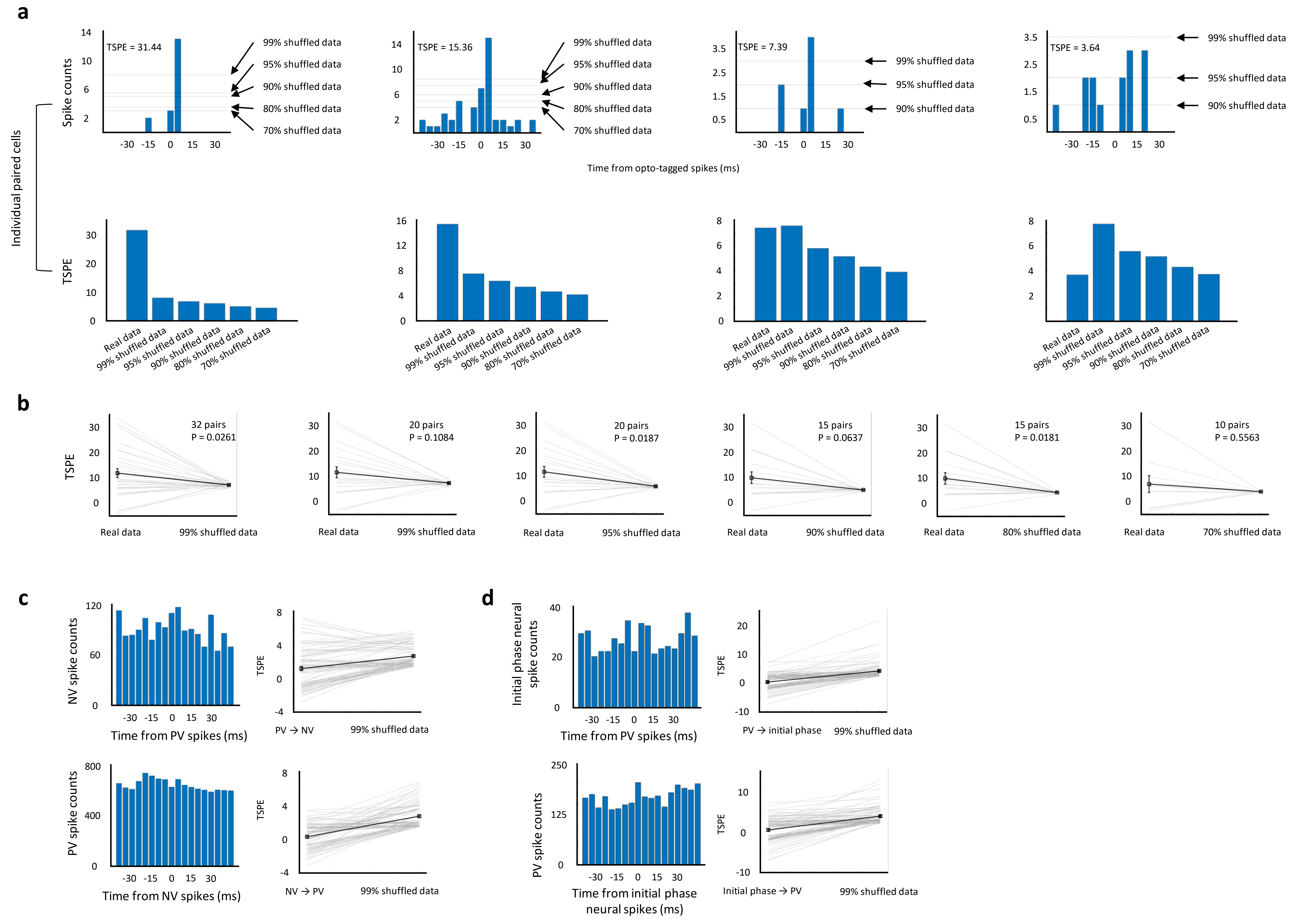


**Supplementary Fig 5. The connectivity among specific neural representation groups**

**a-b.** Connectivity of neural pairs between photo-tagged and network-excited single-units. A: Cross-correlograms of individual neural pairs. B: TSPE comparison of real data and the shuffled data at different percentile. P values are measured by Wilcoxon signed-rank test. **c-d.** Connectivity among classified neural representations. **Left:** Cross-correlograms of individual neural pairs. **Right:** TSPE comparison of real data and the shuffled data at 99 percentile. All mean of real TSPEs are less than mean of 99% shuffled data.

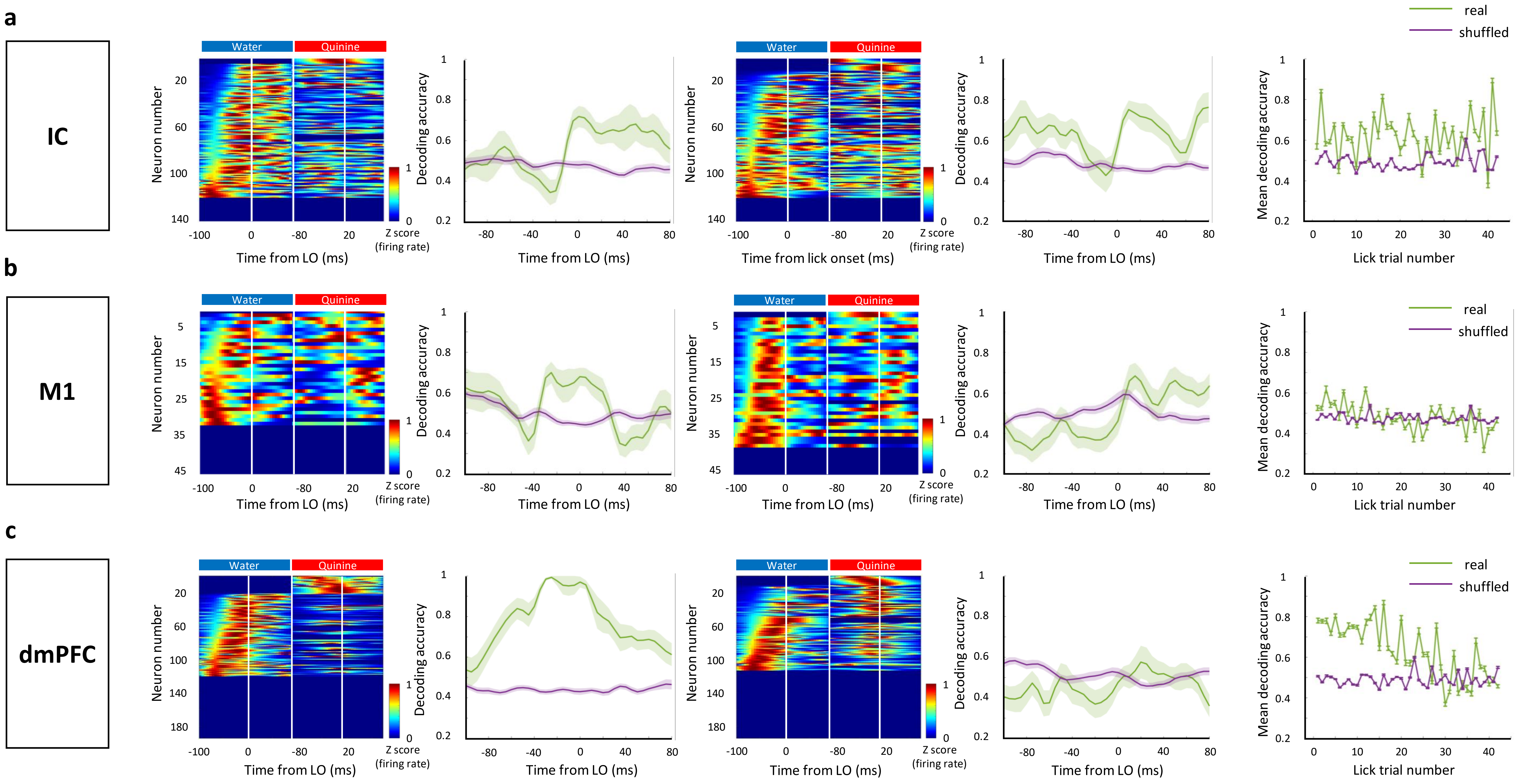


**Supplementary Fig 6. The decoding of liquid types in IC, M1, and dmPFC along with lick proceeding**

**a & b & c. From left to right:** 1$^{st}$ column, color-coded plot showing z scored neural response in the first lick window of 1$^{st}$ recording epoch and 2$^{nd}$ recording epoch; 2$^{nd}$ column, decoding of the first water-lick in 1$^{st}$ recording epoch and in 2$^{nd}$ recording epoch; 3$^{rd}$ column, color-coded plot showing neural response in the 42$^{th}$ lick window of 1$^{st}$ recording epoch and in the 1$^{st}$ lick window of 2$^{nd}$ recording epoch; 4$^{th}$ column, decoding of the 42$^{th}$ water-lick in 1$^{st}$ recording epoch and the first lick in 2$^{nd}$ recording epoch. 5$^{th}$ column, different brain regions showing different liquid type discrimination and changing the discrimination level (decoding performance) along with lick trials evolving. Decoding of water-licks (from 1$^{st}$ to 42$^{th}$ lick trial) in 1$^{st}$ recording epoch and first lick in 2$^{nd}$ recording epoch. Values are mean ± s.e.m.

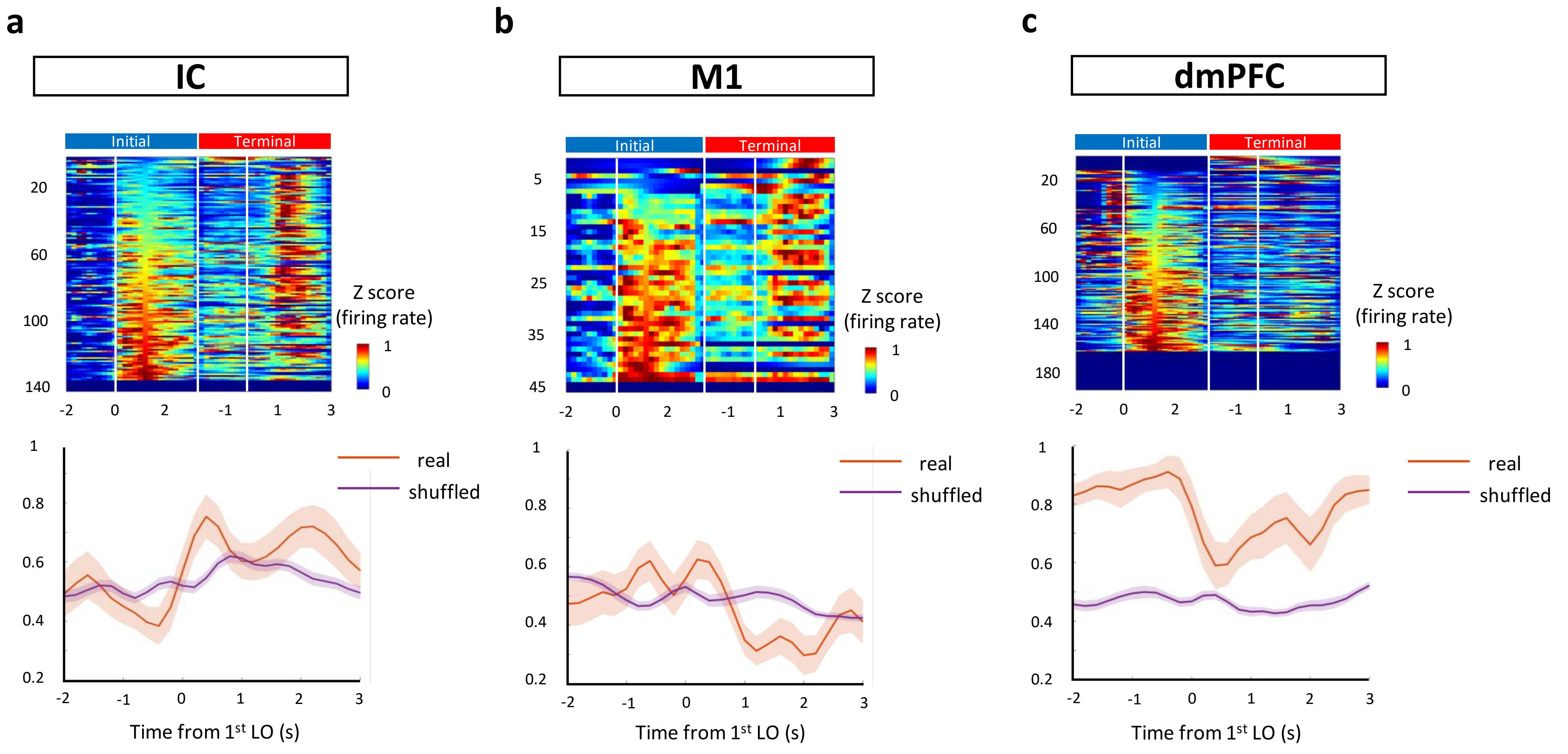


**Supplementary Fig 7. The decoding of initial versus terminal phase in IC, M1, and dmPFC**

**a & b & c.** Decoding of the initial phase in 1st recording epoch (1st water LO-2s to 1st water LO+3s) and the terminal phase in 2nd recording epoch (1st quinine LO-2s to 1st quinine LO+3s). Top: color-coded plot showing z scored neural response in the 1st recording epoch and 2nd recording epoch. Bottom: decoding performance. Values are mean ± s.e.m.

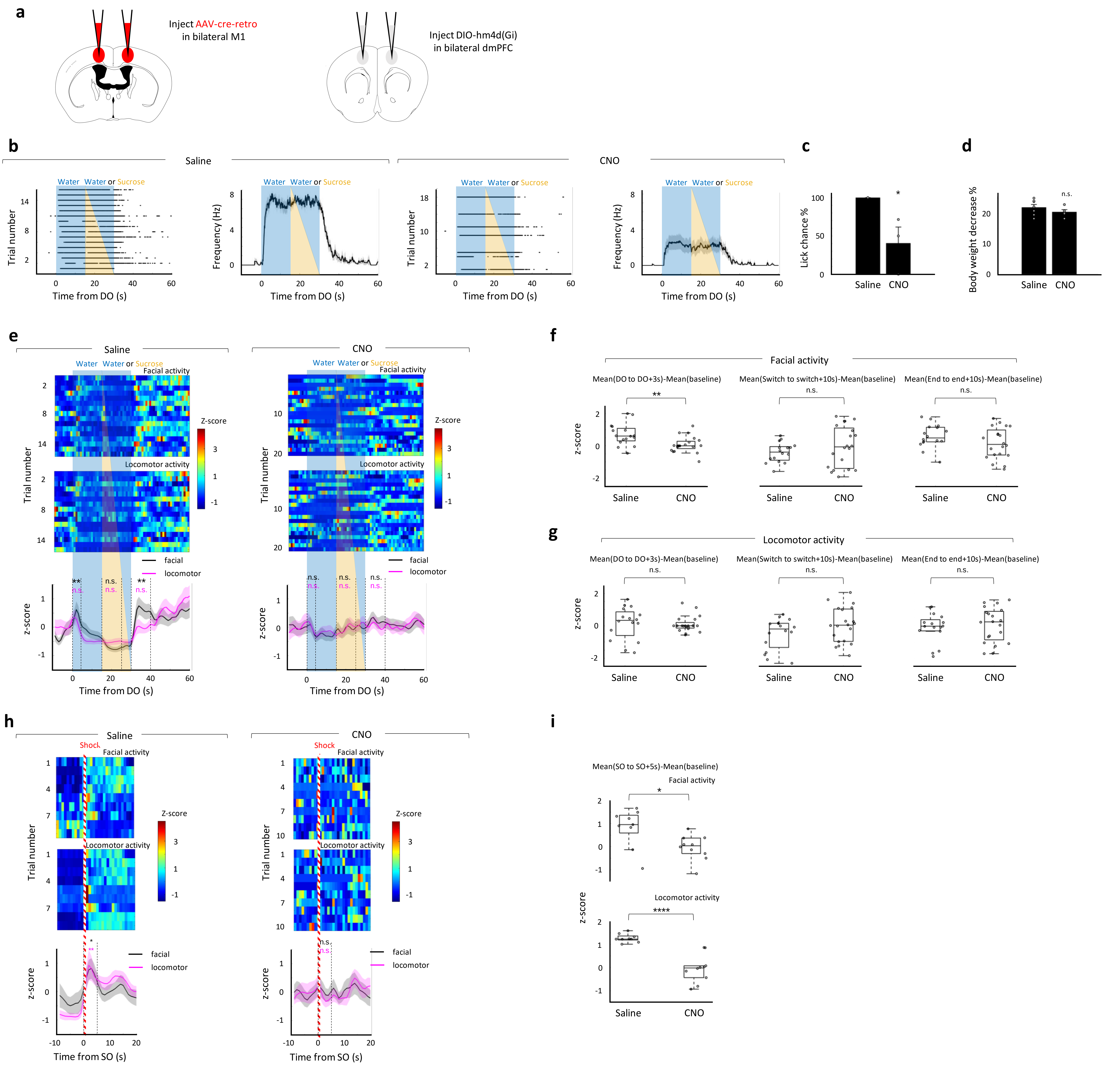


**Supplementary Fig 8. Behavioral effect of dmPFC MP neuron chemogenetically silencing**

**a.** Schematic of bilateral silencing MP neurons in dmPFC. **b.** Lick behavior relative to DO with or without CNO administration. From left to right: 1st column and 3rd column, raster plots showing lick behavior in the session w (15s)-w or s (15s). 2nd column and 4th column, lick frequency plot. **c.** Comparison of lick chance between saline and CNO administrated mice after water delivery. **d.** Comparison of the percentage of body weight decrease between saline (n=16 trials) and CNO (n=20 trials) administrated mice. **e.** Color-coded plot and traces showing z scored facial and locomotor activity relative to water DO. We compared mean facial and locomotor activity in the epochs from DO to LO, from switch onsets to switch onsets+5s, from end onsets to end onsets+5s with that in the baseline. Wilcoxon signed-rank test: *p<0.05, **p<0.01, n.s. not significant. **f-g.** Comparison of z scored facial activity (f) and locomotor activity (g) between saline and CNO administrated mice at various epochs as indicated. n_saline=16; n_CNO=20. Two sample t-test: **p<0.01, n.s. not significant. **h.** Color-coded plot and traces showing facial and locomotor activity relative to shock onset (SO). The mean facial and locomotor activity in the epoch from SO to SO+5s were compared with that in the baseline. Wilcoxon signed-rank test: *p<0.05, ****p<0.0001, n.s. not significant. **i.** Comparison of z scored facial activity (top) and locomotor activity (bottom) between saline and CNO administrated mice at the epoch SO to SO+5s. The trial number is the same as (f-g). For all boxplot, the minima, maxima, and center bounds of box denote 25 percentile, median, and 75 percentile of data, respectively. The upper bound of whisker denote the highest data point, which lower than the sum of maxima bound and 1.5 times of box length. The lower bound of whisker denote the lowest data point, which higher than the subtraction of minima bound and 1.5 times of box length. Statistics: two-sided two sample t-test: *p<0.05, ****p<0.0001.

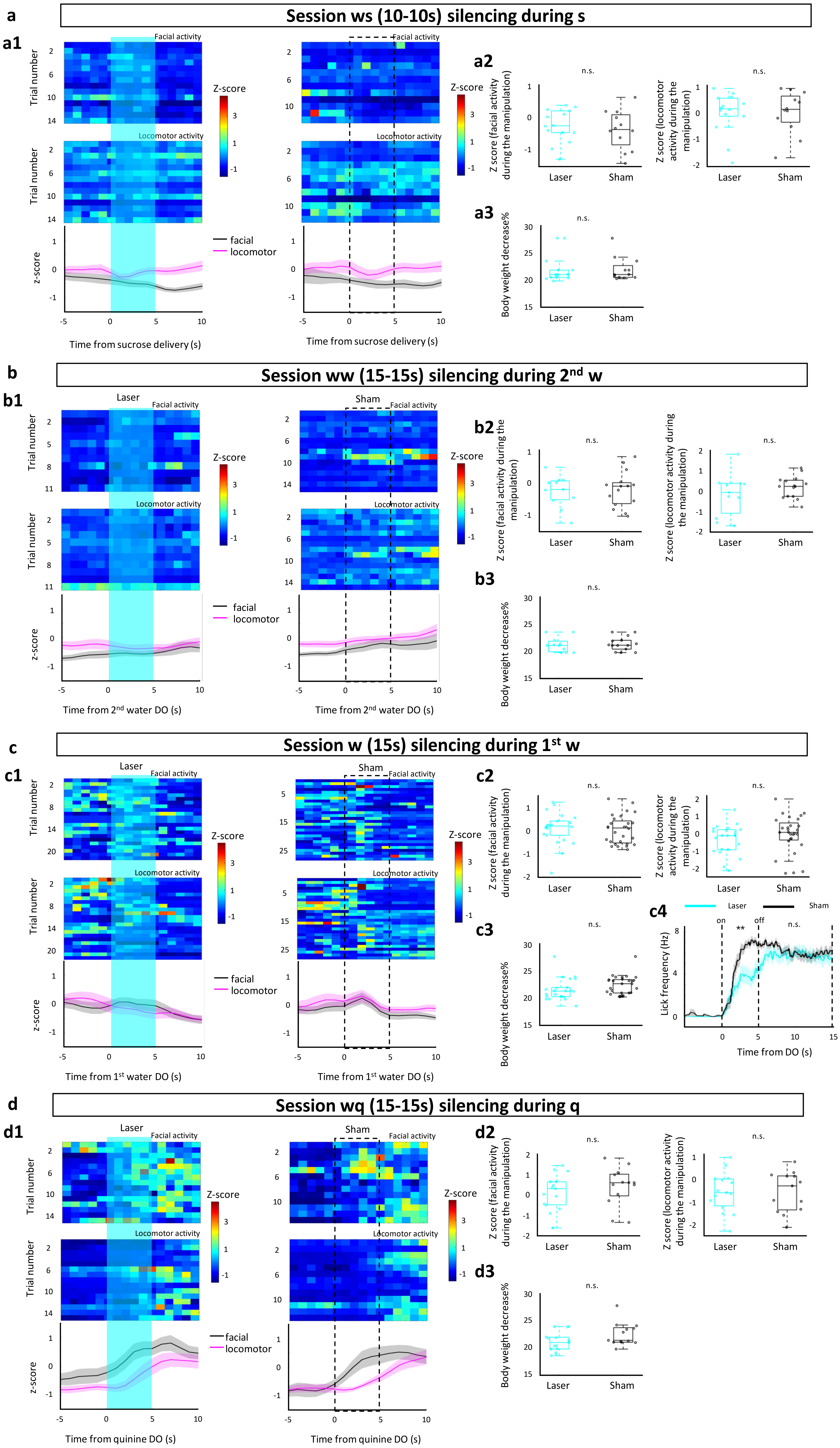


*Legends see next page*

**Supplementary Fig 9. Behavioral effect of dmPFC MP neuron optogenetically silencing**

**a1 & b1 & c1 & d1.** Color-coded plot and traces showing facial and locomotor activity relative to DO. Laser (left) or sham (right) was triggered by DO. Cyan background and dash box represent laser and sham delivery period, respectively. **a2 & b2 & c2 & d2.** Comparison of facial (left) and locomotor (right) activity between laser and sham treatment periods. **a3 & b3 & c3 & d3**. Comparison of the percentage of body weight decrease between laser and sham treatment groups. **c4.** lick frequency relative to DO. The analyzing window was extended to full time period. No significant difference between laser and sham treatment was found after laser or sham off. **a.** n_laser=14; n_sham=12. **b.** n_laser=11; n_sham=15. **c.** n_laser=22; n_sham=28. **d.** n_laser=15; n_sham=13. For all boxplot, the minima, maxima, and center bounds of box denote 25 percentile, median, and 75 percentile of data, respectively. The upper bound of whisker denote the highest data point, which lower than the sum of maxima bound and 1.5 times of box length. The lower bound of whisker denote the lowest data point, which higher than the subtraction of minima bound and 1.5 times of box length. Statistics: two-sided two sample t-test: n.s. not significant **p<0.01

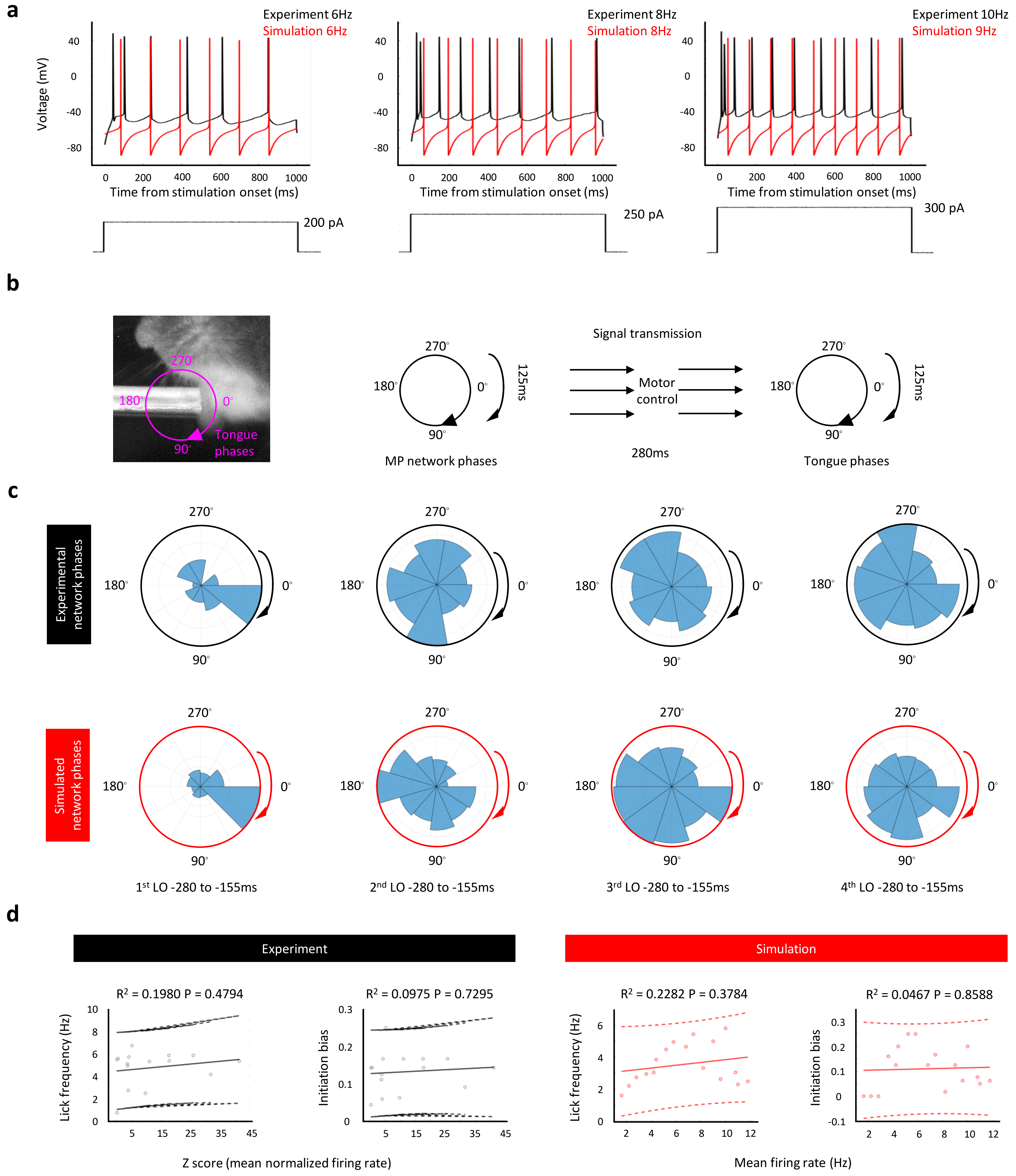


**Supplementary Fig 10. Comparison of the MP network based model and the experimental data**
**a.** Voltage responses of a representative MP neuron in the experiment (data from our previous work[4], black traces) and the model (red traces) to step current injections. **b.** Tongue movement and MP network is quantified by rotational phases. Left: representative image showing tongue movement. Right: one cycle of MP network and tongue movement was set as 125ms and there was a 280ms delay before the signal from MP network arrived to tongue. **c.** Polar histogram of experimental (top) and simulated (bottom) network phases for indicated LO numbers. The blue area of each phase indicates the relative spike counts. **d.** Neural activity of real (left, black) and simulated (right, red) MP network related to lick frequency and initiation bias. Dash lines denote the 95% intervals. R-squares and P values of the linear regression are labeled at each panel. In both **c** and **d**, the experimental data were only chosen from the neurons that the normalized firing rate lower than 50 (65% of the total neuron number) given that slow spiking MP neurons are mainly functional linking deep brain regions and motor cortex[4].

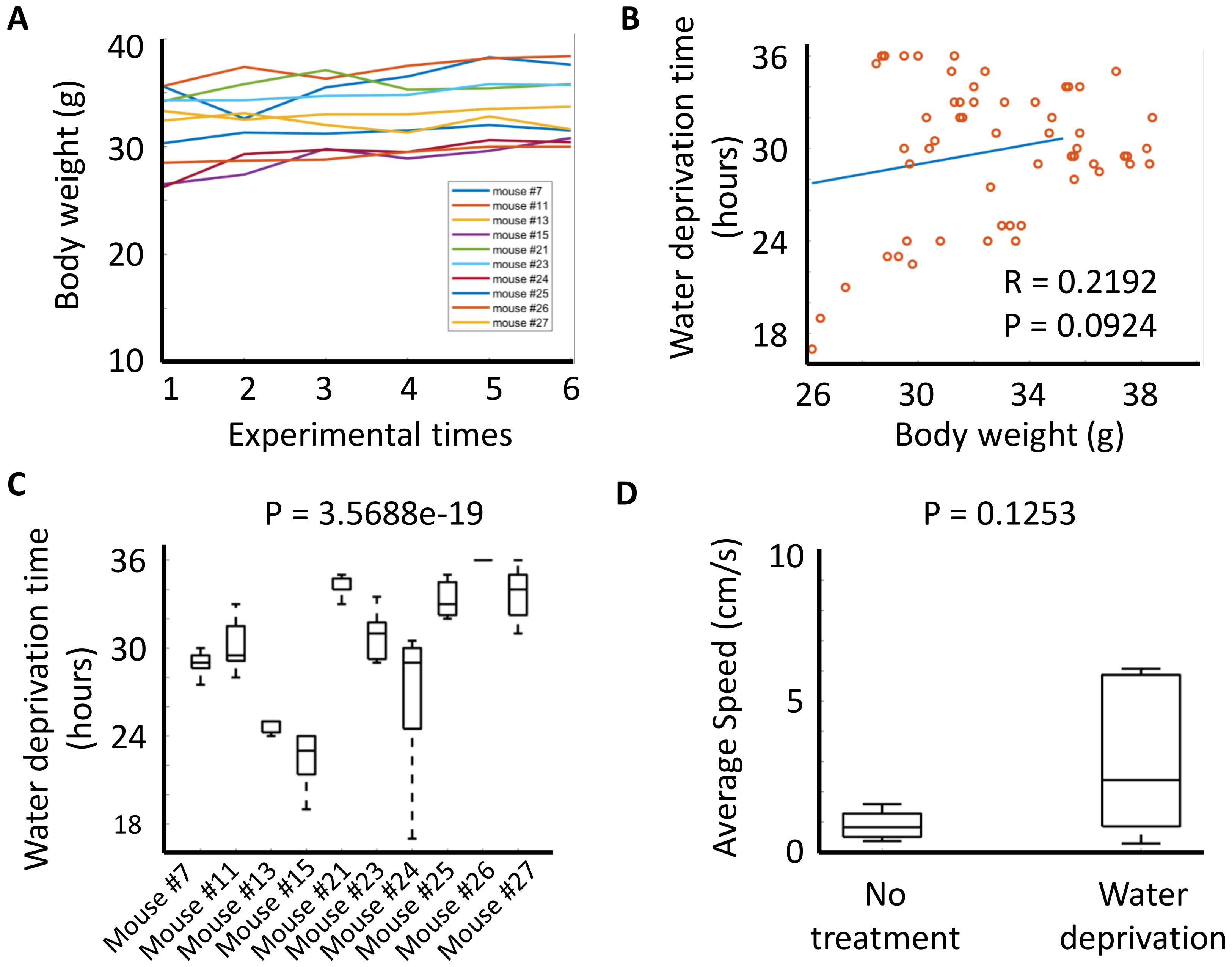


**Method figure 1. A.** Mice body weights before water deprivation across experimental times. The lines with different colors represent different mouse. **B.** Correlation of the body weights before water deprivation and the 22% body weight loss required water deprivation time. **C.** Comparison of 22% body weight loss required water deprivation time across mice. One-way ANOVA, F=30.453, df=9, p=3.5688e-19. n=7. **D.** Comparison of average locomotor activity (represented by average speed) between the mice with and without water deprivation. Statistics: two-sided two-sample t test, t=-15936, df=22, p=0.1253. For no treatment, n=10; for water deprivation, n=12. For all boxplot, the minima, maxima, and center bounds of box denote 25 percentile, median, and 75 percentile of data, respectively. The upper bound of whisker denote the highest data point, which lower than the sum of maxima bound and 1.5 times of box length. The lower bound of whisker denote the lowest data point, which higher than the subtraction of minima bound and 1.5 times of box length.
